# Slow AMPA receptors in hippocampal principal cells

**DOI:** 10.1101/2020.08.05.238667

**Authors:** Niccolò Paolo Pampaloni, Irene Riva, Anna Carbone, Andrew J. R. Plested

## Abstract

Glutamate receptor ion channels such as the α-amino-3-hydroxy-5-methyl-4-isoxazolepropionic acid (AMPA) receptor mediate the majority of fast excitatory neurotransmission in the vertebrate CNS. AMPA receptors canonically provide the fast, millisecond component of the synaptic current. However, we found that about two-thirds of principal cells in the mouse hippocampus express AMPA receptors that do not desensitize and stay active for up to half a second. These receptors are expressed at synapses with a sparse but flat spatial distribution. The resulting increase in charge transfer allows single connections to reliably trigger action potentials. Biophysical and pharmacological observations imply that slow AMPA receptors incorporate γ-8 and other auxiliary proteins, and their activation lengthens individual miniature synaptic currents. Synaptic connections harboring slow AMPARs should have unique roles in hippocampal function.

## Introduction

Neurons in the vertebrate brain receive excitatory input from their presynaptic partners through the rapid activation of glutamate receptors. The fastest of these glutamate receptor ion channels, the AMPA receptors, activate and deactivate in milliseconds in response to glutamate released from synapses(Hestrin et al., 1990). Other types of glutamate receptor show activation over seconds (Gantz et al., 2020; Kidd and Isaac, 1999; Misra et al., 2000), but are either tonically blocked or expressed in only a handful of neurons. Desensitization of native AMPA receptors is near-complete within tens of milliseconds (Colquhoun et al., 1992; Geiger et al., 1995). These properties allow AMPA receptor activation to follow synaptic input with high temporal accuracy, and to participate in short-term depression(Rothman et al., 2009). AMPA channels are retained at synapses by forming complexes with their auxiliary proteins (Bats et al., 2007). These auxiliary subunits act as anchors but also alter receptor responses to glutamate (Tomita et al., 2005). Despite two decades of research, the relevance for brain function of a change in AMPA receptor activity due to auxiliary proteins is lacking.

Auxiliary subunits can slow AMPA receptor kinetics in heterologous expression(Priel et al., 2005) and reduce their tendency to desensitize whilst boosting activation(Coombs et al., 2017). Evidence for desensitization-resistant AMPA receptors in the brain is so far limited to high-frequency activity at certain connections in the cerebellum (Lu et al., 2017; DiGregorio et al., 2007), where hints of a distinct pharmacology were reported (Devi et al., 2016). A mode of enhanced receptor gating with a high conductance (Zhang et al., 2014) driven by activity (Carbone and Plested, 2016) led us to hypothesize that auxiliary subunits inevitably endow AMPA receptors with slow, desensitization-resistant responses. This effect should be particularly prevalent for the γ-8 subunit that is strongly expressed in the hippocampus (Rouach et al., 2005; Yamasaki et al., 2016). Repeated application of glutamate to AMPA-Rs in complex with γ-8 at 10-25 Hz produces a substantial, indefatigable ‘pedestal’ current response (Carbone and Plested, 2016).

To replicate this repetitive activation in the hippocampus, whilst avoiding the potentially confounding effects of presynaptic plasticity, we performed glutamate uncaging at 10 Hz at visually-identified dendritic spines in CA1 pyramidal neurons (Figure 1A). This stimulation reflects a frequency of synaptic input that hippocampal cells might naturally experience during mu or theta waves (Buzsáki, 2002; Takillah et al., 2017). To limit contamination of the responses to glutamate by other receptors/channels, we performed experiments in a cocktail of inhibitors to block GABA-A, kainate, mGluR and GABA-B receptors. We paid particular attention to blocking all types of NMDA receptors, which represent canonical slow glutamate receptor ion channels. To this end, we worked in normal magnesium (2 mM), and included potent NMDA receptor blockers in both intracellular and extracellular media (MK-801 and APV, respectively).

**Fig. 1.**
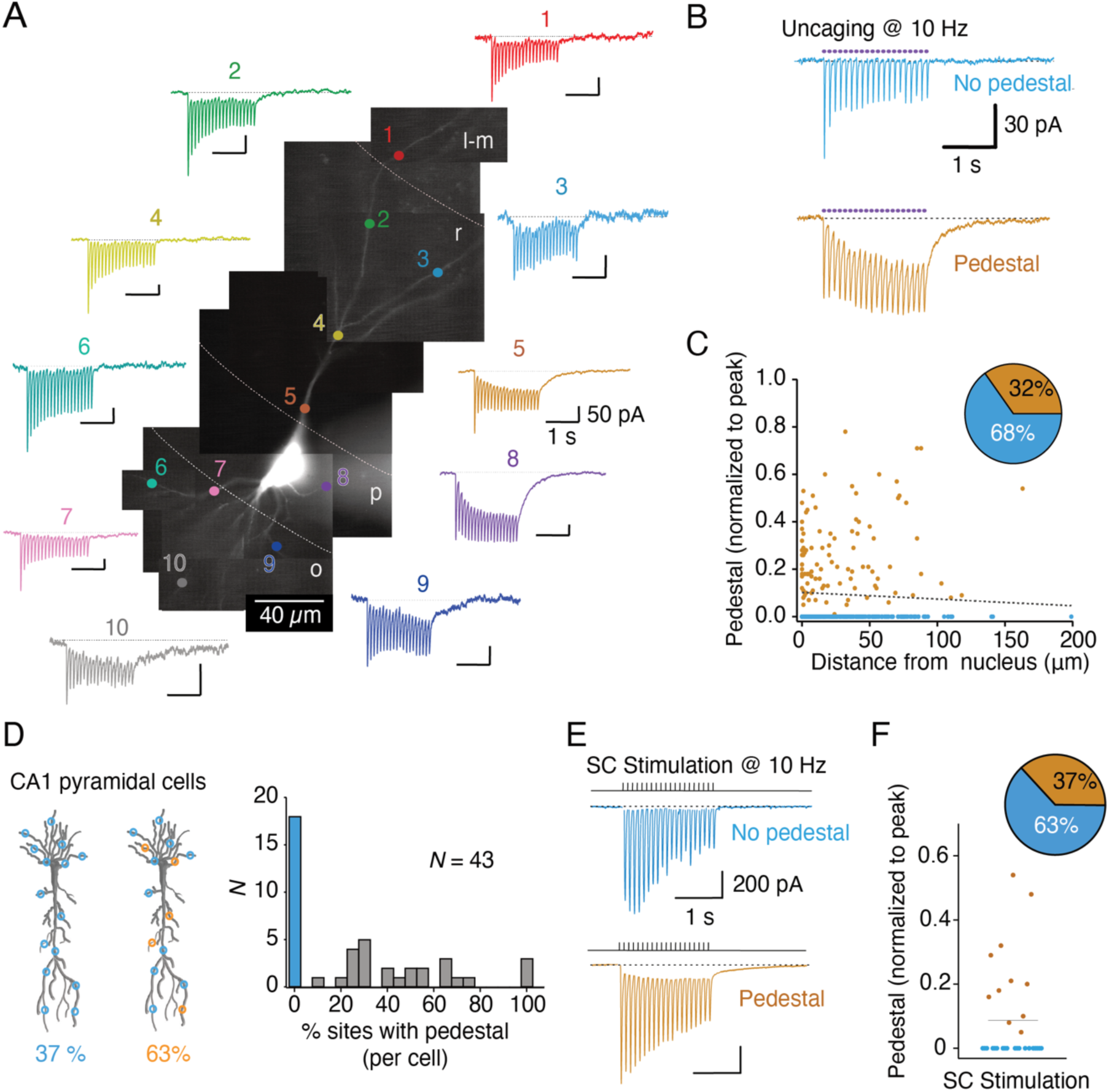
Slow pedestal responses in CA1 pyramidal cells. (A) Tiled fluorescence micrograph of a CA1 pyramidal cell with voltage-clamp responses from 10 Hz uncaging at 10 sites. (B) Examples of typical classical responses (‘no pedestal’) and ‘pedestal’ responses with uncaging pulses indicated as purple circles. (C) Spatial distribution of normalized pedestal (steady-state) current at the end of 20 uncaging pulses at 10 Hz, against distance of the site from the nucleus (dashed line *r*^2^ = 0.003, *n* =285 sites, 43 cells). (D) Incidence and prevalence of pedestal currents. (E) Pedestal and non-pedestal responses generated from 10 Hz Schaffer collateral (SC) stimulation. (F) Distribution of pedestal magnitudes in SC stimulation experiments.

Quite unexpectedly, when uncaging glutamate at 10 Hz, we found both classical fast AMPA receptor responses at some connections and mixed, slow responses at others (Fig. 1). We detected slow pedestal currents in 2/3 of CA1 pyramidal cells, with no relation to distal/proximal dendritic location (Fig. 1) or the peak amplitude of the evoked response (Supplementary Fig. 1). Their distribution appeared to be static, on the timescale of our recordings (10-40 min). The incidence and amplitude of the slow responses showed variability within individual neurons (Fig. 1A), suggesting variable expression of an AMPA receptor with atypical properties. Uncaging experiments in the CA3 region of the hippocampus identified similar but much less prevalent pedestal responses resistant to kainate receptor antagonist UBP 310, in addition to the typical slow kainate receptor currents from the mossy fiber-pyramidal cell synapse (Supplementary Fig. 2)(Castillo et al., 1997; Vignes and Collingridge, 1997).

Could these slow currents be an artifact of glutamate uncaging, recording techniques or variability in our preparation? Several observations suggest this is not the case. First, slow currents were detected in the same cell, and during the same recording, as canonical fast responses. The slow currents present without any treatment or delay following the start of the recording. Pedestal currents were distributed over the neuron with no obvious spatial pattern and were thus dotted along dendrites, interspersed with classical responses (Supplementary Fig. 3). Also, about 1/3 cells from the same slices lacked pedestal responses altogether (Fig. 1D). We also wondered if organotypic culture was a factor in generating large pedestal responses. However, recordings from CA1 pyramidal cells in mouse acute hippocampal slices revealed a similar incidence, form and kinetics of the pedestal current (Supplementary Fig. 4), ruling out this possibility. Finally, we found that the pedestal was retained (with similar kinetics) as we increased the temperature to 32°C, or as we liberated less glutamate by shortening the duration of the UV uncaging pulses (Supplementary Fig. 5). These observations show that any glutamate substantially activating fast, classical responses will also activate slow pedestal currents. Pedestal currents do not require excessive liberation of glutamate, and are slow principally because of slow receptor kinetics.

Glutamate uncaging bypasses presynaptic release, and so we sought to identify pedestal responses according to physiological synaptic function. Recording EPSCs in CA1 pyramidal cells following electrical stimulation of the Schaffer collateral (SC) is one of the most commonly performed experiments in cellular neuroscience. We hypothesized that the pedestal currents should also appear in such recordings, even though their intensity might be different because this experiment entails presynaptic plasticity across multiple connections. A similar proportion of neurons developed a pedestal current when SC electrical stimulation was employed (Fig.1E) to the fraction of neurons showing pedestal responses in uncaging experiments. Importantly, profound glutamate release was not needed to generate the pedestal, because minimal stimulation (with a 10-20% proportion of failures) could still generate a pedestal response when averaging across traces (Supplementary Fig. 6).

Having established the wide incidence of slow, pedestal responses, we sought to better understand their properties. For amplitude-matched responses from the same cell, pedestal responses displayed a much slower decay (550 ± 110 ms) at the end of the 10 Hz stimulation train than their canonical counterparts (40 ± 9 ms; Fig. 2A), which corresponded reasonably well to classical AMPA receptor EPSCs. The very slow response after a train was complemented by a slower decay in response to individual uncaging stimuli (Fig. 2B), whereas the rise time of the response was unchanged. Although distinct from typical fast AMPA receptor activations, these slower decays dovetail well with the slow kinetics of superactive modes of the AMPA receptor (Carbone and Plested, 2016).

**Fig. 2.**
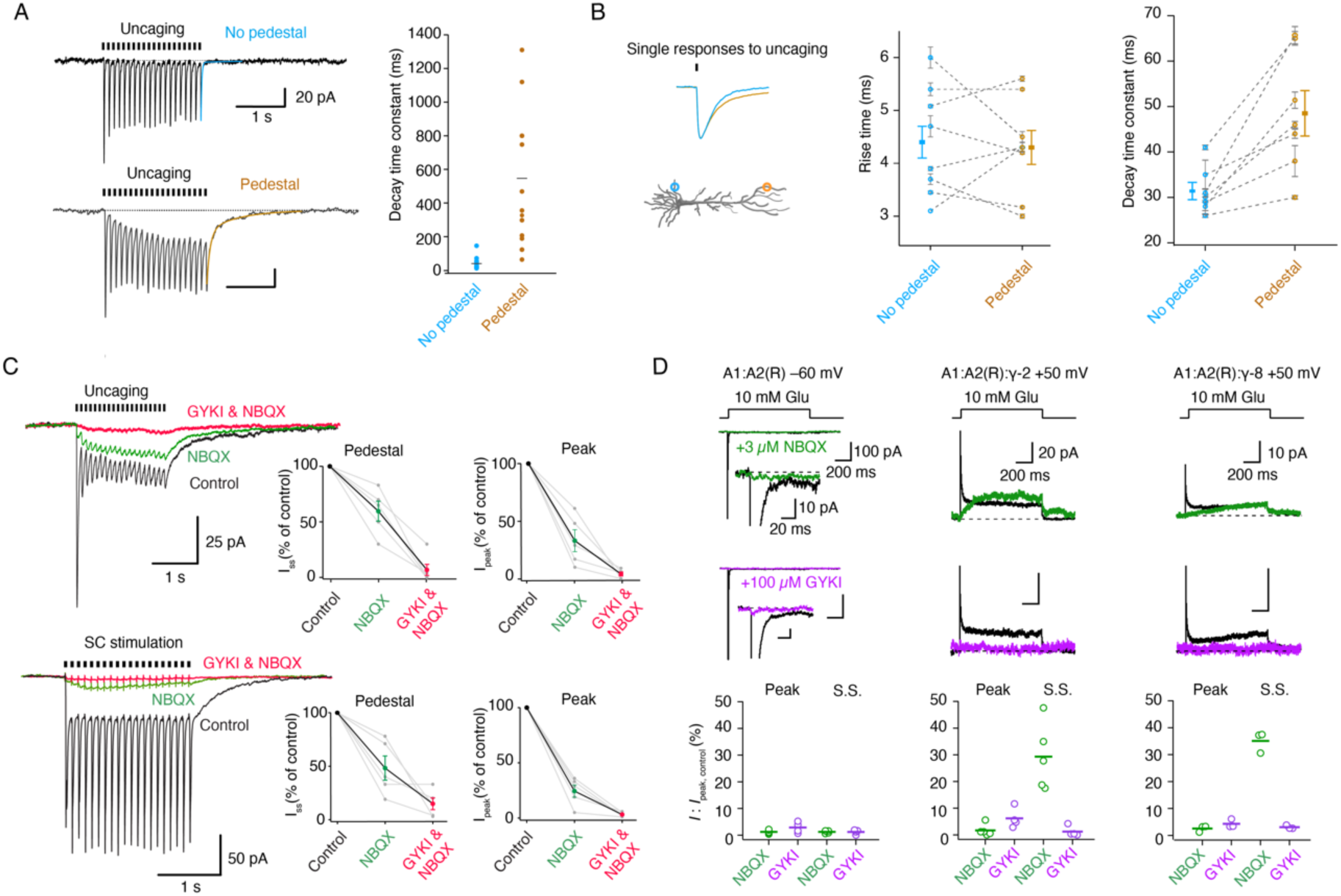
Biophysical and pharmacological properties of pedestal responses. (A) A very slow tail current follows 10 Hz uncaging at pedestal sites whereas responses at ‘no pedestal’ sites have canonical fast decay kinetics. (B) The decays of responses to single uncaging events are slower at identified sites that can generate pedestal currents, but rise times are indistinguishable. (C) In both uncaging and electrical stimulation experiments, pedestal responses are selectively spared by NBQX (1 µM) but abolished by GYKI-52466 (100 µM). (D) Auxiliary proteins γ-2 and γ-8 each endow heteromeric AMPA receptors expressed in HEK-293 cells with pedestal responses resistant to NBQX (3 µM) but not GYKI (100 µM).

Quinoxaline dione antagonists act as partial agonists on AMPA receptors endowed with auxiliary proteins (Menuz et al., 2007). NBQX, likely the most commonly employed AMPAR antagonist(Sheardown et al., 1990), is less effective on receptors in complex with γ-2 (MacLean et al., 2014; Devi et al., 2016). In uncaging experiments, we first confirmed that classical responses were fully blocked by NBQX (not shown, *n* = 5). However, at pedestal sites, NBQX (10 µM) only inhibited the pedestal current by 40% whilst still almost abolishing the fast peak component (Fig. 2C). To confirm that the pedestal currents actually derive from AMPA receptors, we added the AMPA receptor selective non-competitive antagonist GYKI 52466 (Donevan and Rogawski, 1993) at the end of the recording, which entirely blocked the pedestal response. We repeated these experiments with electrical stimulation, and found largely indistinguishable results (Fig. 2C). Consistent with less glutamate being in competition following axonal stimulation than in uncaging, only a lower concentration of NBQX (1 µM) could spare the pedestal generated by electrical stimulation. Pedestal responses were also resistant to N-acetyl-spermine (Supplementary Fig. 7), suggesting the AMPA receptor complexes that generate them contain GluA2 and thus are unlikely to flux substantial calcium.

What could the composition of an AMPA receptor with this unexpected pharmacology be? To answer this question, we performed fast perfusion electrophysiology experiments on defined combinations of AMPA receptor subunits in heterologous expression. In these experiments, we used conditions that tend to isolate receptors enriched with TARPs (Carbone and Plested, 2016). A1A2R plus γ-2 or γ-8 gave similar mix of canonical fast and slow pedestal responses. NBQX (3 µM) selectively inhibited the fast peak currents, whilst sparing or even boosting the slow pedestal, whereas, mirroring our results in pyramidal cells, GYKI 52466 (100 µM) inhibited both components readily. (Fig. 2D). These results strongly suggest that auxiliary proteins endow AMPA receptors with slow ‘pedestal’ activity and NBQX resistance.

To understand the influence of the pedestal expression on CA1 cell activity, we made current clamp recordings (Fig. 3). Following a 10 Hz train stimulation to determine whether a putative synaptic connection had a pedestal response, we switched to current clamp mode and repeated the stimulation. Comparing between non-pedestal and pedestal sites with matched response amplitudes revealed a massive increase in charge transfer (+146 ± 10%) (Fig. 3). Accordingly, uncaging at these sites in current clamp mode had a high probability of triggering an action potential (Fig. 3), correlated to the magnitude of the pedestal (*r*^2^ = 0.64). The mean frequency of spiking during a 10 Hz train was 32% (*n* = 12 pedestal sites) suggesting strong coupling that should be robust in the face of low release probability. In contrast, uncaging at single sites lacking pedestal almost never evoked an action potential (Fig. 3), as expected. We confirmed the specificity of this coupling by addition of GYKI 52466 at the end of the experiment, which blocked the plateau depolarization and the evocation of action potentials (Fig. 3E).

**Fig. 3.**
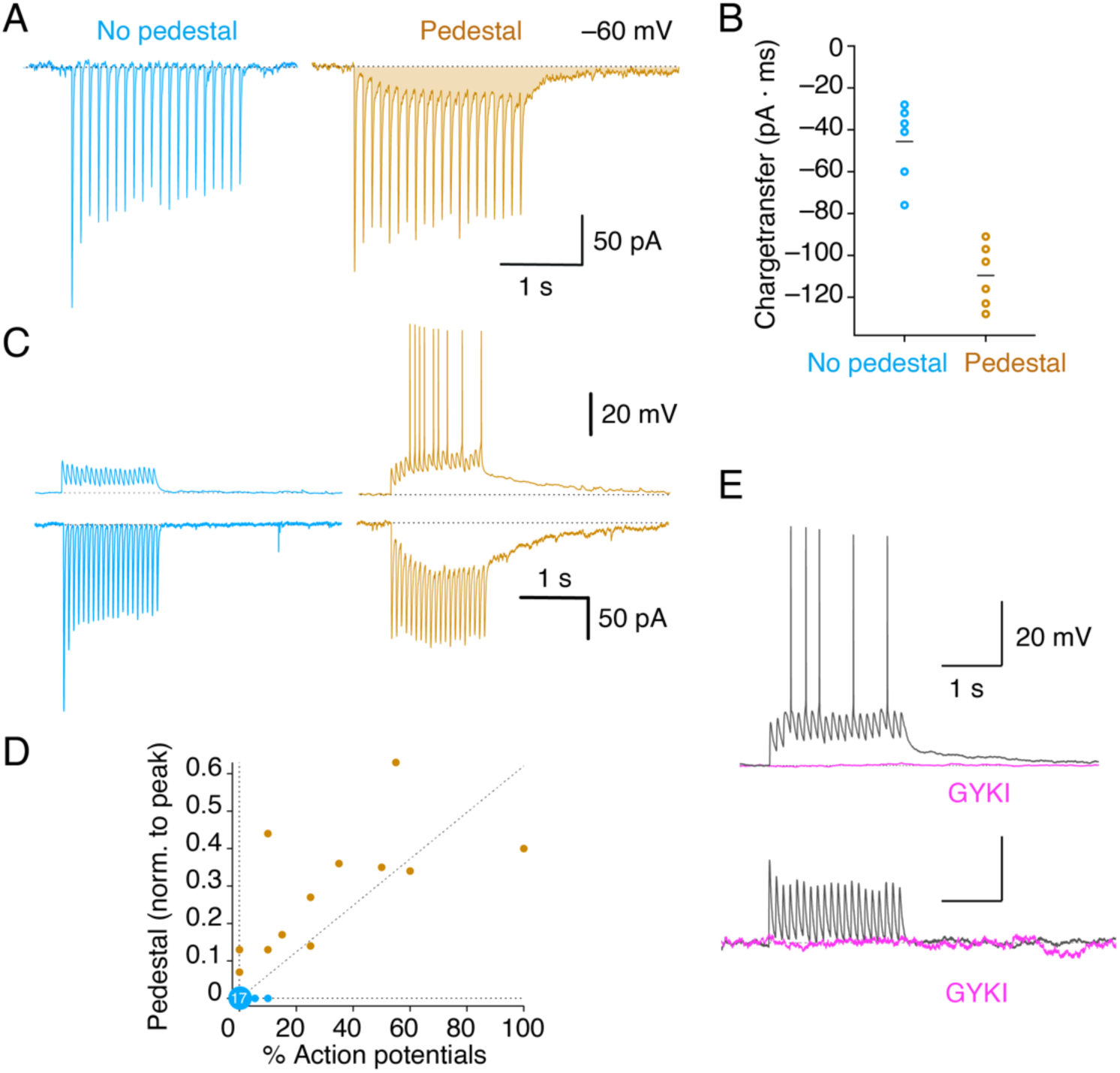
Pedestal connections are instructive, providing a large depolarizing drive. (A) Pedestal responses produce charge transfer of increased amplitude and duration. (B) At amplitude-matched sites in the same cell, pedestal responses produce about 3 times more charge transfer in response to 10 Hz train stimulation. (C) Current clamp recordings show that pedestal sites reliably trigger action potentials whereas similar amplitude canonical responses rarely do. (D) Relation between the pedestal magnitude and reliability of action potential firing during a train. Large blue circle indicates 17 classical sites where zero action potentials were fired from 10 Hz stimulation. Coefficient of determination (*R*^2^) for the line fitted to pedestal responses was 0.64. (E) GYKI-52466 abolished both uncaging EPSPs and the consequent firing of spikes.

A concern from these uncaging experiments might be that glutamate is liberated over a wide volume. Therefore, even though we expect that AMPA receptors are concentrated at synapses, we could conceivably have obtained pedestal currents by activating a special class of receptors that are systematically excluded from synapses. These receptors might therefore avoid native synaptic release. Unlikely as this scenario is, we sought to confirm directly that slow AMPA receptors were located at synapses and activated even by spontaneous release. Although two-photon uncaging is the gold standard for releasing glutamate in the most accurate way, the volume liberated is still large compared to vesicular release, and such experiments would not confirm that synaptically-released glutamate activates slow AMPA receptors.

To address synaptic localization directly, we recorded mEPSCs in naïve neurons for 5 minutes before performing our normal scan of 10-20 putative synaptic sites with glutamate uncaging. This scan allowed us to characterise the prevalence of pedestal responses. Miniature ESPCs were longer in neurons where we could identify pedestal currents (Fig. 4), and the lengthening of the decay beyond 20 ms correlated with the incidence of the pedestal currents (Fig. 4). This experiment shows that receptors with slower kinetics participate in individual synaptic currents, suggesting they have a similar synaptic localization to classical AMPA receptors. Miniature EPSCs were also substantially larger on average in cells that had strong pedestal prevalence (Supplementary Fig. 8). Overexpression of γ-8, the most abundant TARP in the hippocampus (Tomita et al., 2003), by single cell electroporation further lengthened mEPSC decays, without affecting the amplitudes of mEPSCs or evoked responses (Supplementary Fig. 8). Strikingly, overexpression of γ-8 increased the prevalence of pedestal currents in the transfected neurons to ∼100% (Fig 4B and Supplementary Fig. 8), further confirming that pedestal currents and slow mEPSC decays are due to AMPA receptors. Single cell electroporation with an inactive γ-8 mutant (Riva et al., 2017) abolished large pedestal responses but increased the incidence of the pedestal overall (Supplementary Fig. 9), consistent with γ-8 expression not being saturated and other auxiliary proteins being involved in the production of slow AMPA currents. Null γ-8 also reduced the amplitude of synaptic responses, consistent with experiments from γ-8 knockout mice(Rouach et al., 2005), making these results difficult to interpret.

**Fig. 4.**
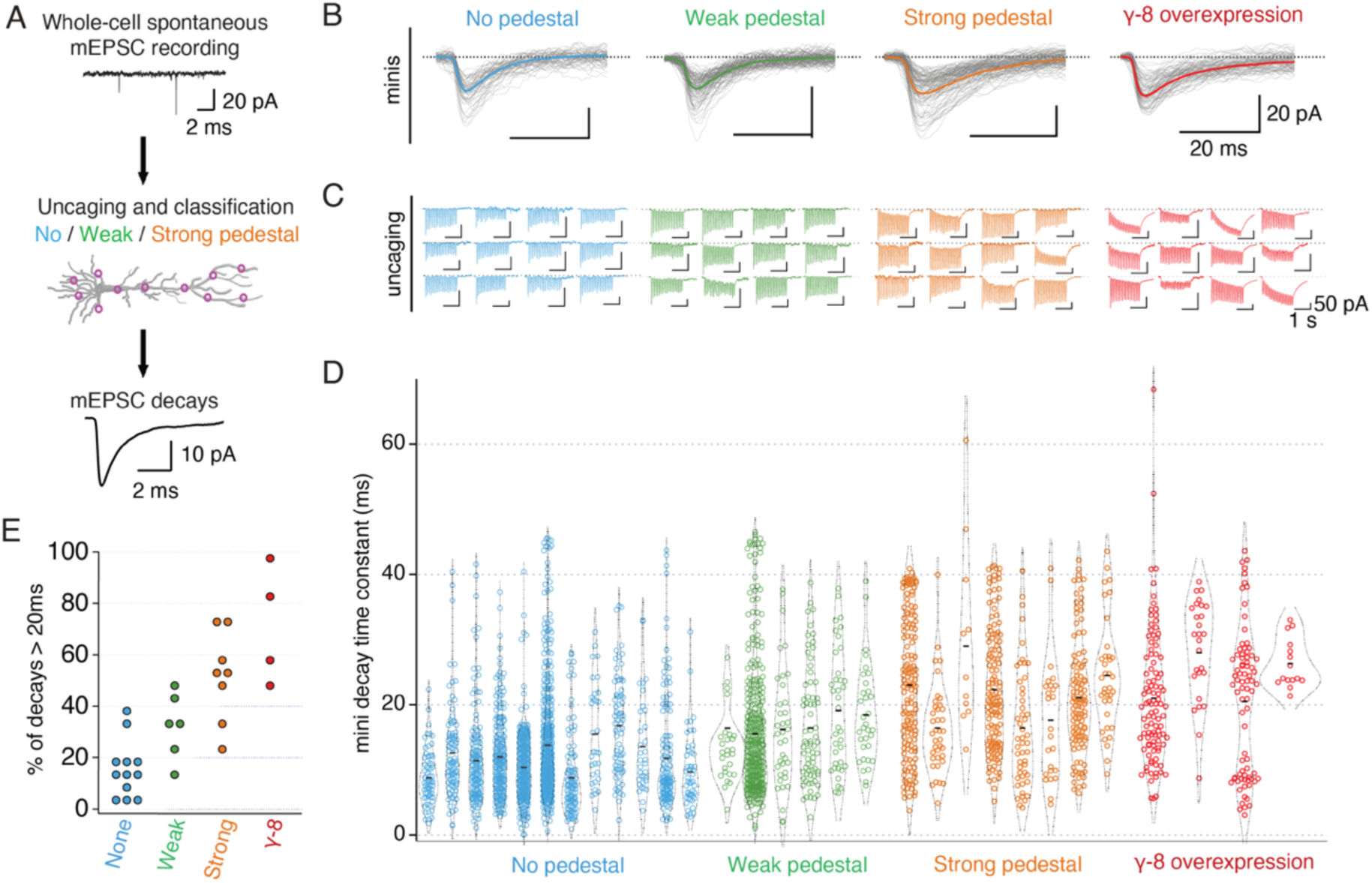
Pedestal currents correspond to slow individual miniature synaptic currents. (A) Experimental design. Miniature currents were recorded for >5 minutes in a naïve cell preceding uncaging survey to classify the pedestal prevalence. (B) Aligned miniature currents from 4 exemplary cells (C) Uncaging surveys from 12 sites for each color-coded cell in panel B. (D) Fitted decay constants of individual miniature currents. Each column represents a cell classified according to the *post hoc* uncaging results. (E) Summary of miniature currents with long decays. Cells without pedestal have a low prevalence of long decaying miniature currents, and increasing incidence of pedestal correlates with lengthening decay.

An obvious question is: if pedestal currents are so prominent, why were they not previously reported? To some extent, the right experiments were not done. More importantly, we fear that investigators have selected synaptic responses that fit best the canonical view of the AMPA receptor as a purely fast ion channel receptor. It is probable that many investigators systematically discarded experiments in which slow responses were observed (for example, CA1 pyramidal cell responses from SC stimulation), which would correspond to 1/3 of cells in our hands.

Our isolation of desensitization-resistant slow AMPA receptors, likely associated with γ-2 or γ-8 subunits, suggests that a subset of extremely powerful excitatory inputs are present in hippocampal CA1 and CA3 pyramidal cells. Previous work suggested that TARPs are ubiquitously expressed in CNS neurons(Tomita et al., 2003), and that TARP modulation of gating is widespread(Menuz et al., 2007). Therefore, slow AMPA currents are likely found elsewhere in the hippocampus and cortex. With more intense stimulation than used here, slow AMPA currents were also reported in cerebellum (Devi et al., 2016; Lu et al., 2017) and calyx (Taschenberger et al., 2002). The apparently higher affinity for glutamate of these receptors in complex with TARPs (Coombs et al., 2017) reduces the absolute requirement for direct apposition to vesicle release sites(Raghavachari and Lisman, 2004) and dilutes the dependence of synaptic currents on receptor diffusion(Heine et al., 2008) and clustering(Savtchenko and Rusakov, 2014). At the same time, pedestal currents are much more potent at depolarizing target neurons than classical fast responses. Our results urge further care in the interpretation of results where quinoxaline dione antagonists are employed to block AMPA receptors(Menuz et al., 2007; Cossart et al., 2002). In terms of short-term synaptic plasticity, classical presynaptic mechanisms of potentiation with repetitive activity due to calcium accumulation(Katz and Miledi, 1968) now find a natural complement in the progressive augmentation of slow AMPA receptor currents. Pedestal currents represent a long-sought short-term potentiation mechanism from a purely postsynaptic locus(Zucker and Regehr, 2002). The implications of instant, long-lasting depolarizations from a single input for long-term plasticity induction and synaptic integration are likely to be substantial(Remy and Spruston, 2007). Slow AMPA currents have implications for homeostatic regulation(Turrigiano, 2008), and the excitability of any cells where they are found, with knock on effects for network function. Finally, it is intriguing to speculate as to whether pedestal currents participate in feature recognition (Bittner et al., 2015) or enable particular inputs to rapidly instruct cellular responses to stimuli or environments, such as conversion into place cells(Epsztein et al., 2011).

## Acknowledgments

We thank Andrew Penn, Mario Carta and Mauro Pulin for advice on slice culture and electroporation; Sabine Fievre and Christoph Mulle for help with initial experiments in CA3 cells; Francesca Logiacco for advice on acute slice preparation; Paul Kammermeier for advice on pharmacology; Mark Mayer, Estelle Toulmé and Christoph Schmidt-Hieber for comments on the manuscript and Gert Rapp and for assistance with SysCon software and microscope optimization. N.P.P. was recipient of an EMBO Long Term Fellowship (no. ALTF 873-2018), and A.L.C. was recipient of a NeuroCure Female Fellowship. This work was funded by the Deutsche Forschungsgemeinschaft (DFG) Heisenberg Professorship (PL619/3-1), DFG RU2518 DynIon (P3; PL619/5-1), the DFG under Germany’s Excellence Strategy – EXC-2049 – 390688087 –NeuroCure and the ERC grant 647895 “GluActive” (all to A.J.R.P.). This manuscript is dedicated to the memory of James R. Howe, whose generous sharing of data before publication was critical at early stages of this work.

## Author contributions

N.P.P. conceived and performed experiments, analysed data and wrote the paper; I.R. conceived and performed experiments and analysed data A.L.C. conceived and performed experiments and A.J.R.P. conceived and performed experiments, analysed data and wrote the paper.

## Methods

### Organotypic Slice culture preparation, single cell electroporation

350 µm-thick organotypic hippocampal slice cultures were prepared from P6 to P9 WT C57 mice. Animals were maintained in compliance with the EU Legislation on the protection of animals used for scientific purposes and were approved by Landesamt für Gesundheit und Sociales Berlin (LaGeSo). Slices were prepared on filter paper according to the interface method (Stoppini et al., 1991; De Simoni and Yu, 2006) and cultured in a MEM-based mouse slice culture medium, with the addition of 15 % Horse Serum; 1x B27; 25 mM HEPES; 3mM L-Glutamine; 2.8 mM CaCl_2_; 1.8 mM MgSO_4_; 0.25 mM Ascorbic Acid; 6.5 g/L D-Glucose. 3 days after plating, the medium was replaced and then exchanged every 4 days. Cultures were grown in an incubator with 5% CO_2_ at 34°C.

Single cell electroporation (SCE) was performed at 15-16 days of slice culture (Wiegert et al., 2017). Visually identified CA1 pyradmidal neurons were transfected by single cell electroporation. Slices were placed in the microscope chamber in the presence of 3-4 ml sterile pre-warmed (34°C) HEPES-based artificial cerebrospinal fluid (aCSF) containing 145 mM NaCl, 2.5 mM KCl, 2 mM CaCl_2_, 1 mM MgCl_2_, 10 mM HEPES and 10 mM glucose, adjusted to 310 mOsm /l and pH 7.3 with NaOH at room temperature. Borosilicate glass pipettes were filled with intracellular solution containing 135 mM K·CH_3_SO_3_, 4 mM NaCl, 2 mM MgCl_2_, 2 mM Na_2_ATP, 0.3 mM Na_2_GTP, 0.06 mM EGTA, 0.01 mM CaCl_2_, 10 mM HEPES, adjusted to 300 mOsm/l and pH 7.2-3.

For SCE experiments, cDNA was added to the intracellular solution the day of the experiment at a concentration of 4 ng/µl. A pipette with resistance in saline of 8-11 MΩ was brought to the cell body and held in loose-cell attached configuration. DNA was delivered into the target neuron through stimulation of 500 ms at 50 Hz with a pulse amplitude of -10 to -12 V, using an Axoporator 800A (Axon Instruments). After transfection, slices were placed back into the incubator, and the culture medium was enriched with 10 μg/ml Gentamicin. Transfected cells were recorded 24 to 48 hours after the electroporation.

### Preparation of acute hippocampal slices

Hippocampal Acute slices were obtained from postnatal day P18–P21 mice, using a standard protocol (Papouin and Haydon, 2018). Briefly, after cervical dislocation, the brain was quickly removed from the skull and placed in ice-cold slicing solution (aCSF) containing (in mM): 10 Glucose, 125 NaCl, 1.25 NaH_2_PO_4_, 2.5 KCl, 26 NaHCO_3_, 2 MgCl_2_, 1 CaCl_2_, saturated with 95% O_2_ and 5% CO_2_, pH 7.3–7.4. Transverse hippocampal slices (300 µm thick) were cut with a Leica VT 1200S vibratome and stored at room temperature in a holding bath containing the same solution as above. After incubation for at least 1 h, an individual slice was submerged in the recording chamber and continuously superfused at a rate of 5 ml/min with oxygenated experimental ACSF containing (in mM): 10 Glucose, 125 NaCl, 1.25 NaH_2_PO_4_, 2.5 KCl, 26 NaHCO_3_, 1 MgCl_2_, 2 CaCl_2_, saturated with 95% O2 and 5% CO2, pH 7.3–7.4.

### Electrophysiology and glutamate uncaging in organotypic hippocampal cultures and acute slices

Somatic whole cell patch clamp recordings of visually identified CA1 principal cells were performed after 16-23 DIV with microelectrodes (3-8 MΩ) prepared from borosilicate glass capillaries (1B150F-4 World Precision Instruments) using a Sutter P-1000 puller. Slices were superfused with recirculating aCSF (5 mL/min at room temperature) containing 4-methoxy-7-nitroindolinyl-glutamate (MNI-caged-L-Glutamate; HelloBio HB0423) at a concentration of 0.5 mM and used for the whole day of recordings.

Glutamate was uncaged by a 405 nm laser (one-photon excitation) mounted on a custom-built upright microscope (Scientifica SliceScope). The collimated beam was directed through a UGA-42 Firefly laser scanner (Rapp Optoelectronic GmbH), and directed through a Zeiss epifluorescence reflector (Examiner A1) into the water dipping 60x objective (Olympus LUMPlanFL N; N.A. 1). The uncaging laser and Jenoptik ProgRes MF camera were controlled SysCon software (Rapp) and ImageJ (https://imagej.nih.gov/ij/) respectively. After reaching the whole cell modality, we waited several minutes to allow the diffusion of AlexaFluor-594 dye (ThermoFisher; 20 μM, dissolved into the intracellular solution) into the processes of the neuron. Afterwards, dendritic ROIs were selected by illuminating the sample with a 595 nm LED (Thorlabs). UV light pulses (usually 1 ms) were delivered by triggering the laser. Simultaneous passage of red emission (600 nm) and 405 nm light for uncaging was achieved by using a 405/488/594nm Laser Triple Band filter set (TRF 69902; Chroma) mounted in a Zeiss TIRF cube. At the end of each experiment, we documented the uncaging sites and their distances from the cell nucleus using a tiled, multifocal plane fluorescence micrograph of the filled neuron, neglecting the z-displacements which were typically small.

For voltage clamp recordings neurons were held at –60 mV (not corrected for the liquid junction potential which was calculated to be –6.6 mV). To monitor the uncompensated series resistance (<20 MΩ), a hyperpolarizing voltage step (–10 mV, 100 ms) was delivered at the beginning of each recording. Recordings with series resistance changes >20% were discarded. For the current clamp experiment, the bridge balance compensation was adjusted before the beginning of each recording. Most recordings were done at room temperature (23°C)(Herring et al., 2013), but for some recordings we heated the bath inflow with a TC02 inline heater (Multichannel Systems, Germany), adjusting as needed to produce a final temperature of 32°C in the bath.

To evoke whole cell EPSCs in CA1 principal cells from stimulating the Schaffer Collateral, we used an electrode that had resistance of 1-3 MΩ when filled with aCSF. The stimulating electrode was placed in CA3/CA1 interface at the stratum radiatum to activate Schaffer collateral/commissural afferents (300-500 µm away from the recording electrode). Monopolar stimulation was applied with an Iso-Flex constant-current stimulator (API Instruments, Jerusalem, Israel), and the stimulation trigger was controlled by Axograph software, (20 × 1 ms pulses at 10 Hz).

For most experiments in slices, drugs were added to the aCSF at the following concentrations: AP-5 (HelloBio; 20 μM), SR-99531 (HelloBio; 10 μM), CGP-55845 (Tocris; 10 μM), (RS)-MCGP (Tocris; 200 μM) and UBP-310 (HelloBio; 10 μM), while MK-801-maleate (HelloBio; 1mM) was added to the intracellular solution. For current clamp recordings the same pharmacological cocktail was used, but we omitted TTX. For the experiments involving stimulation of the Schaffer Collateral, besides the exclusion of TTX from the ACSF, QX-314 (5 mM, HelloBio) was added to the pipette solution.

Data were acquired with a Multiclamp 700B amplifier (Molecular Devices) and digitized at 20 kHz under the control of Axograph (Axograph Scientific).

### HEK 293 cell culture and electrophysiology

Human embryonic kidney (HEK) 293 cells were co-transfected with the AMPAR subunits GluA1 and GluA2 (edited at the Q/R site) plus the accessory protein TARP γ2 or γ8. γ2 was transfected using a DNA mass ratio of 1:1:2 GluA1:A2:γ2, whereas for γ8 the ratio was 1:1:5 GluA1:A2:γ8. Patch-clamp experiments in the outside-out configuration were performed 24 hours after transfection. Cells were selected based on simultaneous EGFP and DsRed fluorescence signals indicating co-expression of GluA2 and the TARP respectively. The presence of GluA1 subunits was assessed based on the current-voltage (I-V) relationships showing partial voltage-dependent block by intracellular spermine. The intracellular solution contained (in mM): 120 NaCl, 10 NaF, 0.5 CaCl_2_, 5 Na_4_BAPTA, 5 HEPES and 0.05 spermine. The extracellular solution contained (in mM): 150 NaCl, 0.1 MgCl_2_, 0.1 CaCl_2_ and 5 HEPES. Both solutions were titrated to pH 7.3. Using a piezo-driven tool for fast application (Physik Instrument, Germany), outside-out patches were held in extracellular control solution and exposed to brief pulses (400-700 ms) of 10 mM glutamate. To test inhibition, either 3 μM NBQX or 100 μM GYKI-52466 were added to both control and glutamate barrels. Currents from AMPAR-TARP complexes were acquired at the holding potential of +50 mV. All recordings were sampled at 10 kHz and filtered with a 5 kHz low-pass filter using an Axopatch 200B amplifier (Molecular Devices, U.S.A.) and AxoGraph. Because both NBQX and GYKI abolished the fast peak response to glutamate, we normalized the effects of NBQX or GYKI-52466 on the peak and steady state currents as the ratio of the peak (*I*_pk, antagonist_) or steady-state (*I*_ss, antagonist_) response to the mean peak current amplitude (*I*_pk, control_) evoked by glutamate in the absence of antagonist.

### Data analysis

Data were analyzed offline with Axograph, Clampfit (Molecular Devices) and Igor 8 (WaveMetrics). For each uncaging site or whole cell evoked current, the pedestal value was calculated as the steady state current (*I*_ss_) in ratio to the peak current after 20 pulses at 10 Hz. The *I*_ss_ was taken to be 0 when smaller than 5 pA. All the recorded traces were low-pass filtered at 1 kHz. Analysis of spontaneous miniature AMPA-mediated synaptic excitatory currents (mEPSCs) was performed using a threshold set at three times the root mean square (RMS) noise. For each recording a template based on the events recorded was then generated (Axograph). This template was then used to collect the mEPSCs, from which we measured their amplitudes and decays (90 to 10 % of the peak). Putative miniature currents with rise times slower than 5 ms were discarded.

For the analysis of the charge transfer, cells with a similar I_peak_ at the 20^th^ pulse (113 ± 4 pA for non-pedestal and 112 ± 3 pA for pedestal, *n* = 6 for both groups) were selected. Similarly, for the firing probability of pedestal vs non-pedestal inputs, inputs were selected so that the evoked amplitude at the 20^th^ pulse was 100 ± 20 pA. For this analysis, only inputs within 70 μm from the cell nucleus were analyzed, in order to minimize spatial filtering effects.

