## Supplementary Figures 1-9 for "Slow AMPA receptors in hippocampal principal cells"

### Supplementary Figures 1 to 9

#### Slow AMPA receptors in hippocampal principal cells

**Niccolò Paolo Pampaloni, Irene Riva, Anna Carbone, Andrew Plested**

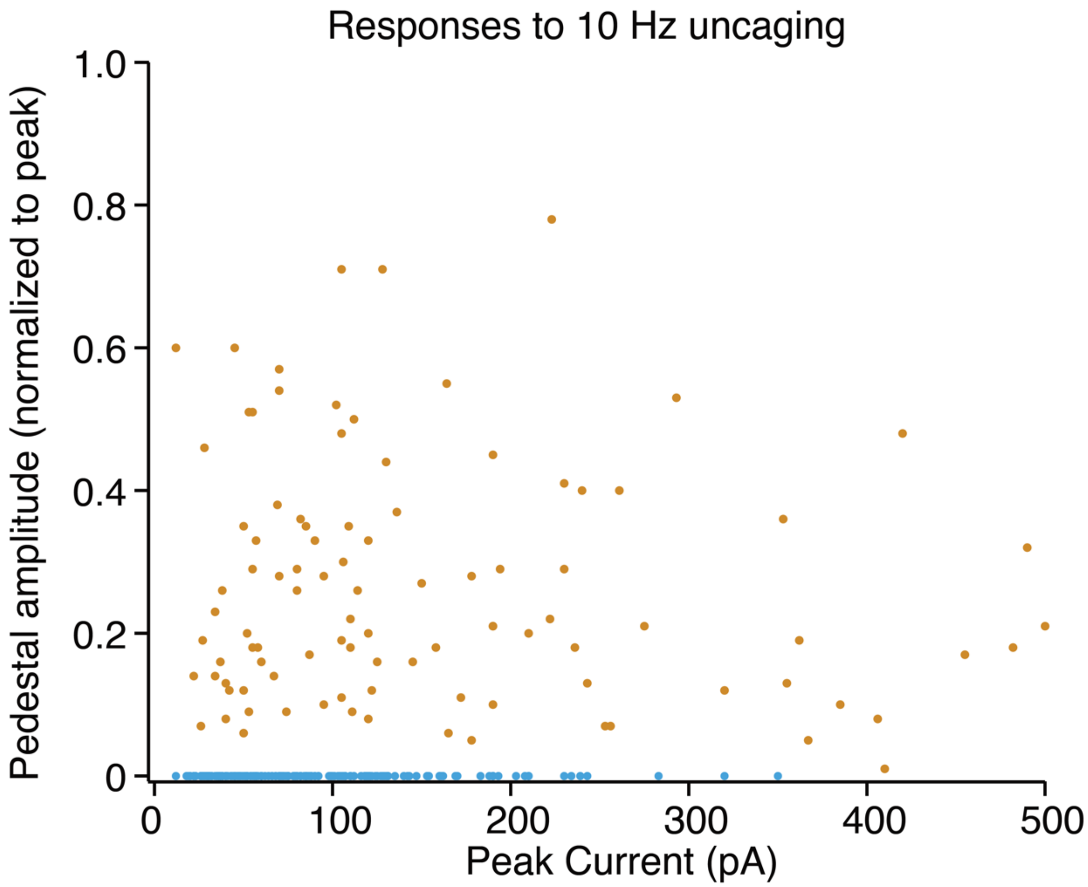


**Supplementary Figure 1. Magnitude of the normalized pedestal against response amplitude.**

The distribution of the normalized pedestal amplitude ($I_{ss}/I_{peak}$) in CA1 pyramidal neurons following 10 Hz uncaging. There is no relation between the magnitude of the pedestal response and the response amplitude, for 285 uncaging trials from 43 cells.

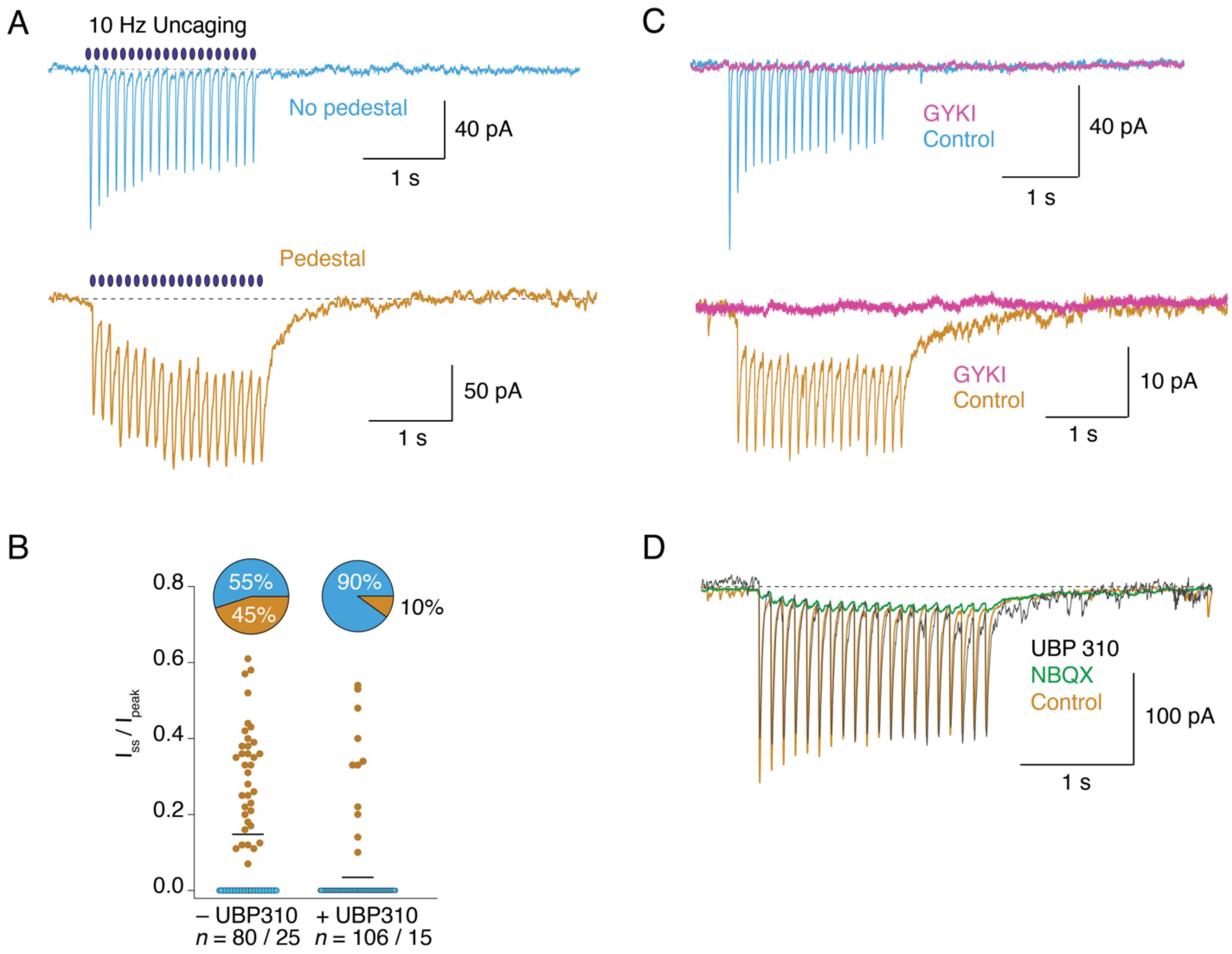


**Supplementary Figure 2. Pedestal current incidence and pharmacology in CA3 pyramidal cells**

(A) Pedestal responses (orange) and classical fast responses (blue) to 10 Hz uncaging were detectable in CA3 pyramidal cells.(B) The incidence of pedestal currents was less in UBP310 (10 $\mu$M, 10% of uncaging sites), suggesting a strong contribution of kainate receptors to these responses in CA3 neurons. Number of uncaging sites and cells are indicated. (C) Classical fast and slow pedestal currents were completely blocked by the AMPA selective antagonist GYKI 52466 (100 $\mu$M), (D) At a site showing resistance to UBP310 (20 $\mu$M), the pedestal component was selectively resistant to NBQX (1 $\mu$M, green).

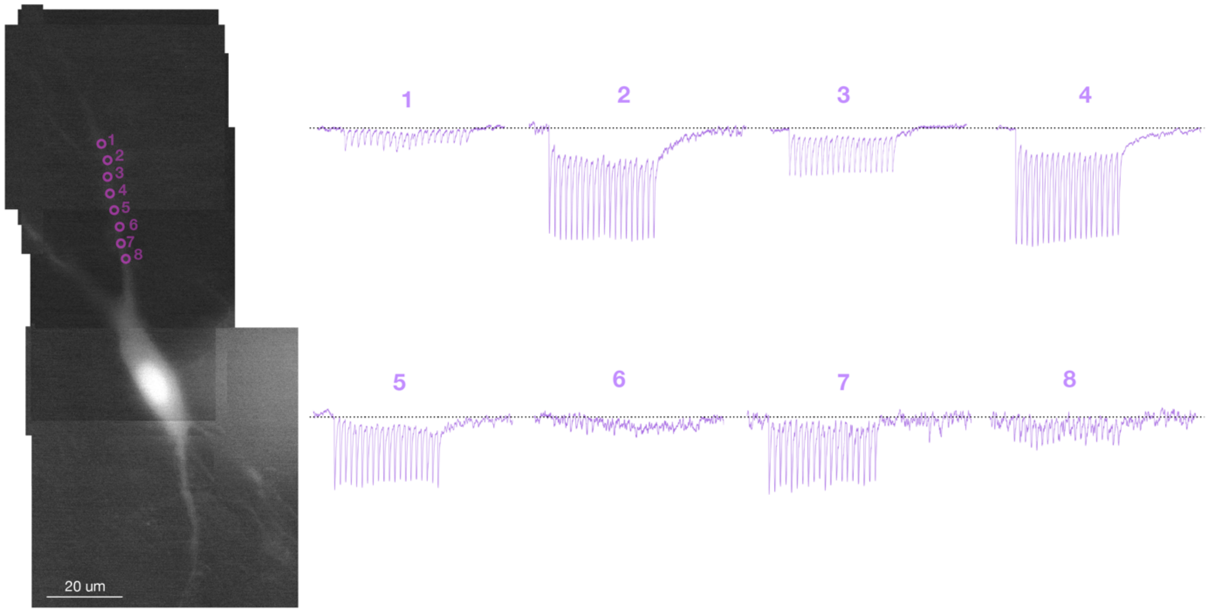


**Supplementary Figure 3. Walking along the dendrite with 5 $\mu$m pitch**

Uncaging at 8 sites along a CA1 pyramidal cell dendrite in a single whole cell patch clamp recording reveals both pedestal and classical responses interleaved with non-responsive sections of shaft. Neuron filled with Alexa 594 from the patch pipette.

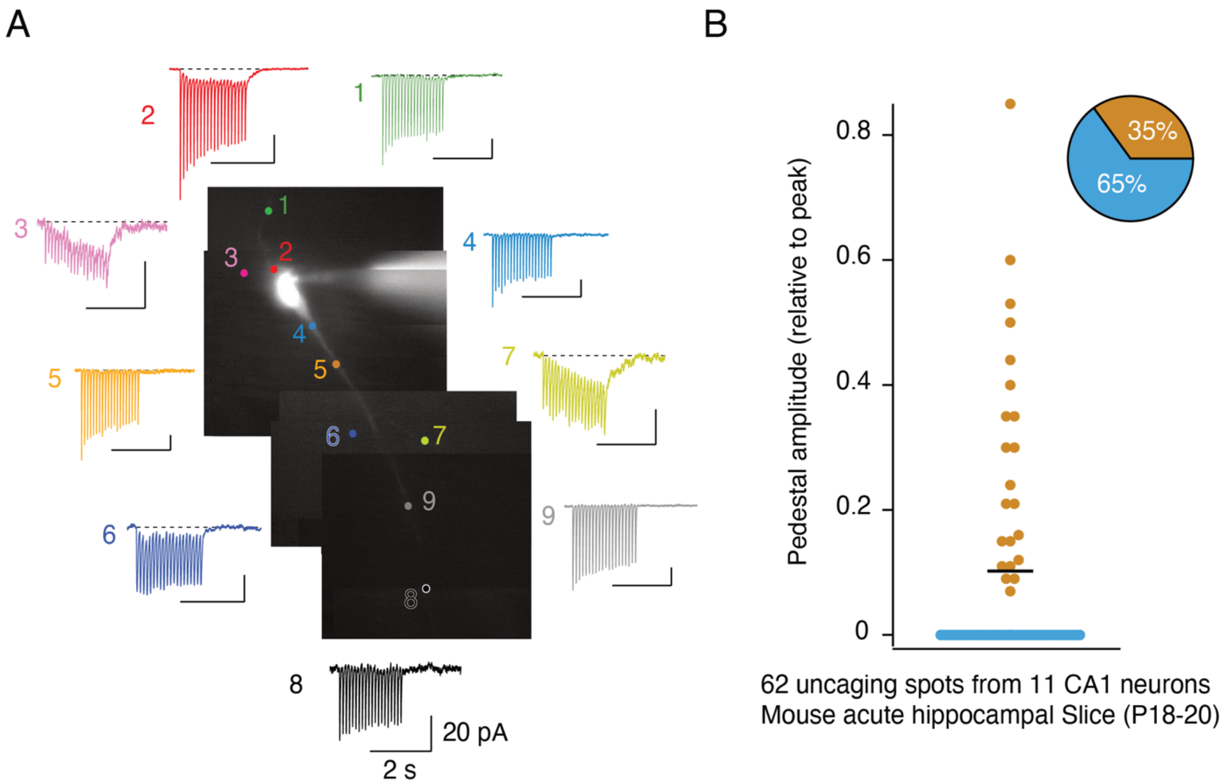


**Supplementary Figure 4. Pedestal prevalence and magnitude in acute mouse hippocampal slice**

(A) Responses to 10 Hz uncaging at 9 sites in a CA1 pyramidal neuron. A slow pedestal response is evoked at sites 2, 3, 6 and 7. Neuron filled with Alexa 594 from patch pipette (at soma). (B) Summary of uncaging at 62 sites in 11 pyramidal cells from acute slices (P18-20 mice), showing similar incidence of pedestal responses (orange, 35% of sites) to recordings made from mouse organotypic hippocampal cultures.

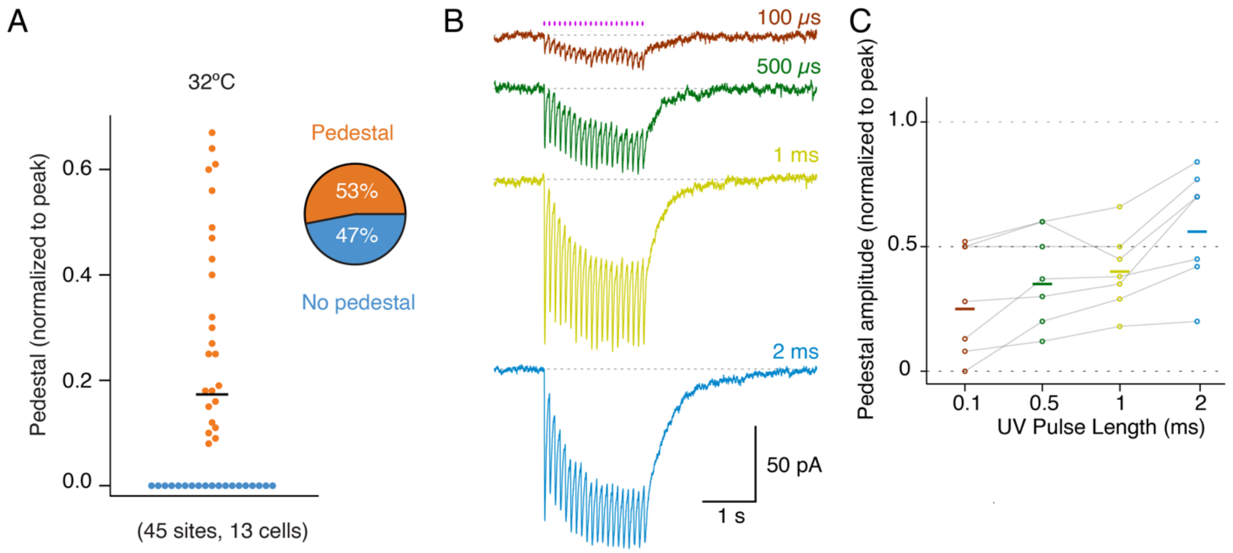


**Supplementary Figure 5. Negligible effects of changing temperature and UV pulse length.**

(A) Recordings made in organotypic slices at 32ºC have a similar incidence and magnitude of pedestal response to those at room temperature. Kinetics of the decay at the end of a 20 pulse, 10 Hz uncaging train were also indistinguishable between room temperature at 32ºC (pedestal decay 550 ± 100 ms, $n = 14$ responses, and non-pedestal decay 32 ± 4 ms, $n = 15$ responses). (B) Typical pedestal responses to 10 Hz uncaging at the same site, colour coded by pulse duration. The magnitude of both the fast response and the slow pedestal is proportional to the pulse duration. The pedestal is present even with 100 $\mu s$ pulses of 405 nm laser (brown trace). (C) The ratio of pedestal amplitude to peak increases about 2-fold across a 20-fold range of pulse lengths.

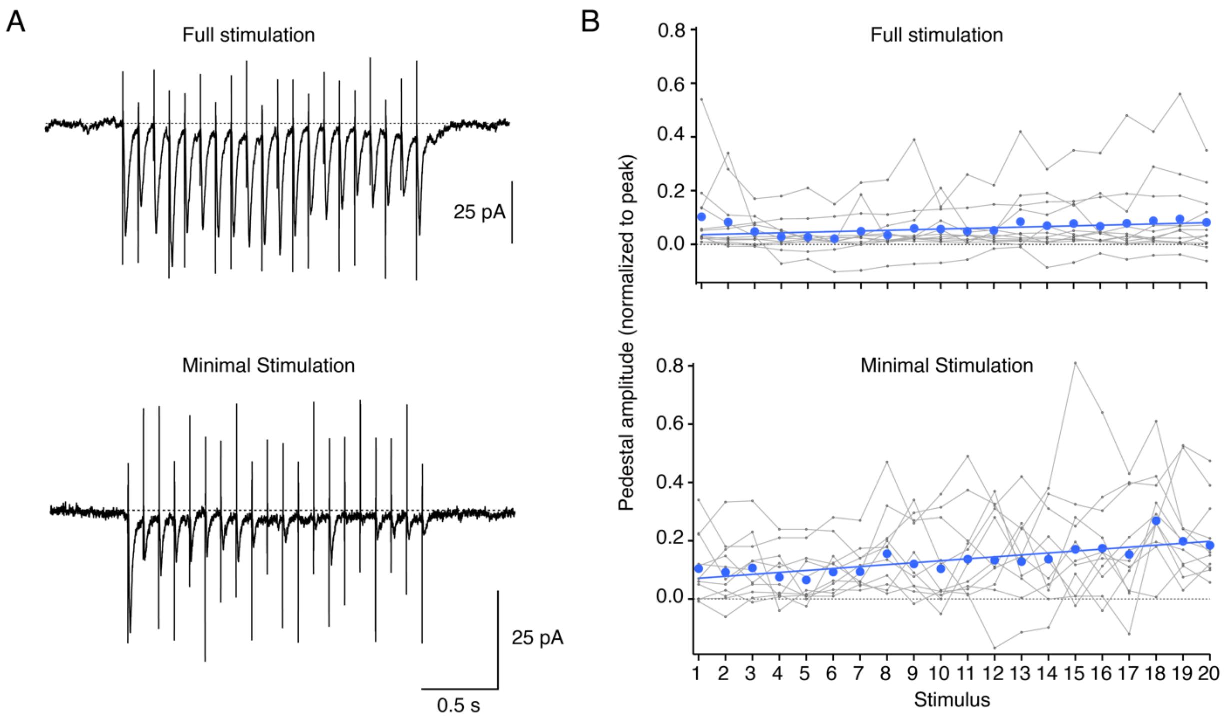


**Supplementary Figure 6. Pedestal responses with minimal versus full stimulation and analysis**

(A) Recording of CA1 pyramidal cells with 10 Hz Schaffer Collateral stimulation to evoke robust responses (*upper panel*) and minimal stimulation (*lower panel*, with ~15% failures) (B) Both stimulation regimes evoke a reliable pedestal response on average (blue line; 12 neurons).

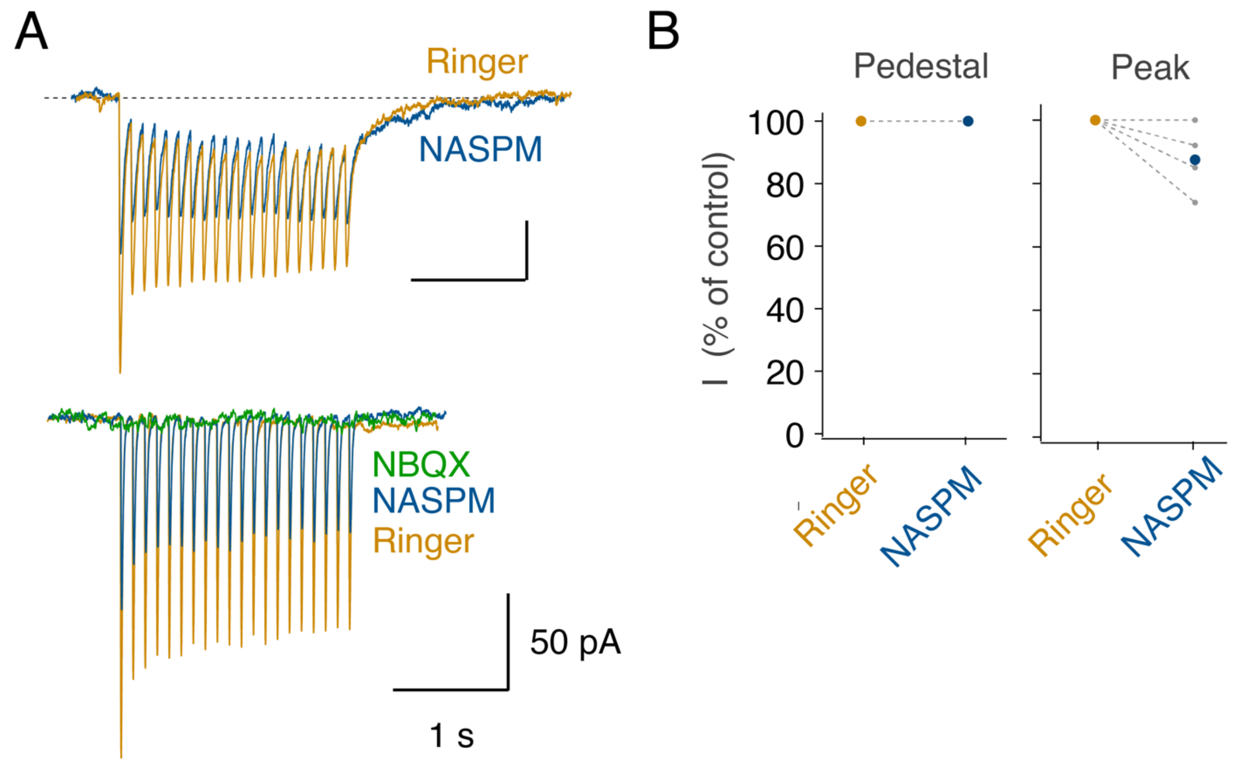


**Supplementary Figure 7 NASPM has no effect on the pedestal**

(A) N-acetyl-spermine (NASPM, an inhibitor of AMPA receptors lacking GluA2) did not inhibit the pedestal current. In an example recording of a pedestal response (*upper panel*), the peak currents (measured from the pedestal level) were inhibited on average about 20% but the pedestal current was untouched. A classical response to 10 Hz uncaging (*lower panel*, without pedestal) was inhibited about 50% by 25 µM NASPM and abolished by NBQX. (B) Summary of results from 5 separate recordings.

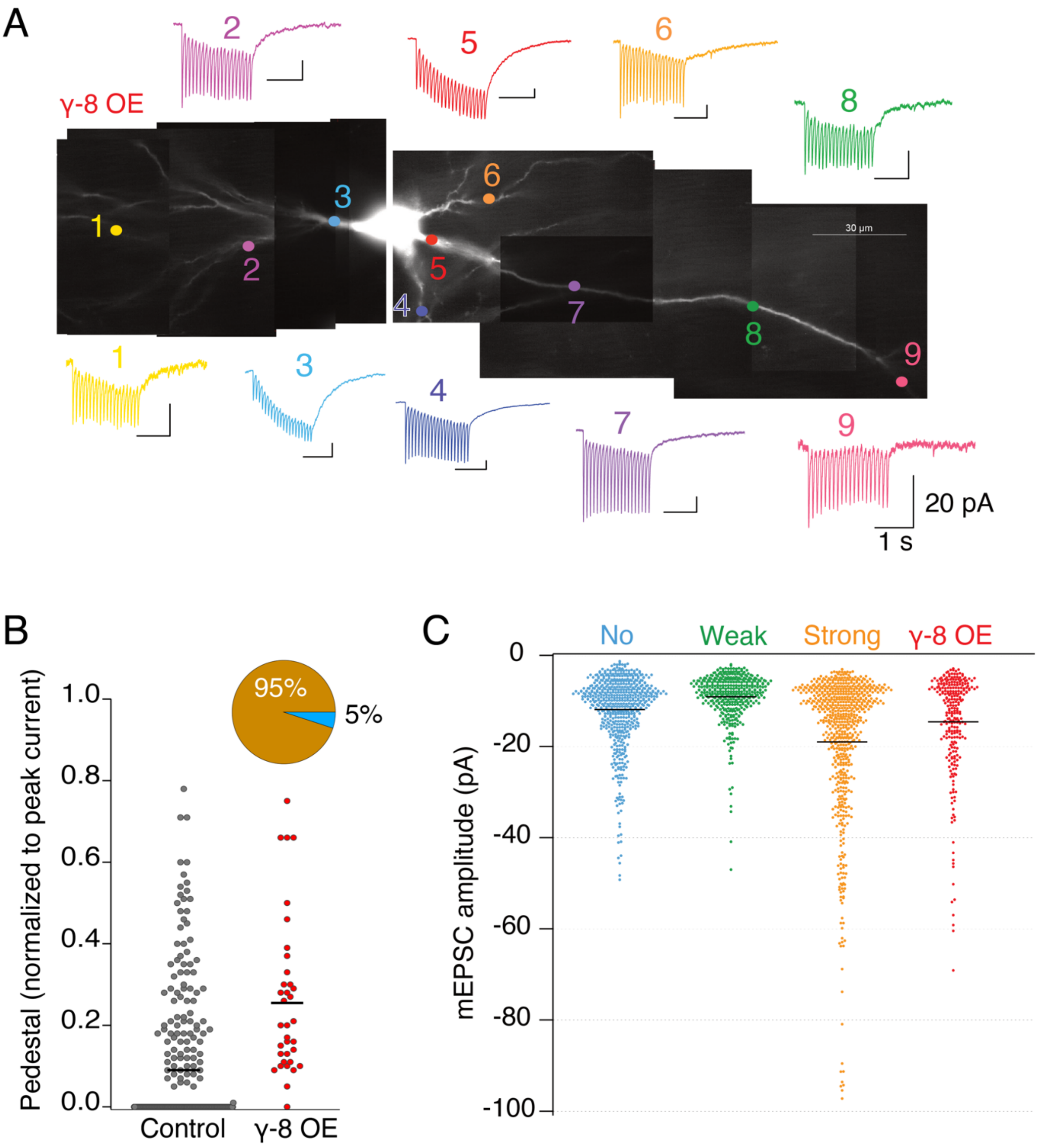


**Supplementary Figure 8. Gamma-8 overexpression, uncaging and miniature current amplitudes**

(A) Uncaging at 10 Hz at 9 sites on a neuron electroporated with the auxiliary subunit gamma-8. All sites show pedestal, although the magnitude of pedestal at site 9 was small. (B) Overexpression of gamma-8 increased the fraction of pedestal responses to 95% ($n$ = 4 cells). (C) Miniature current amplitudes were mildly increased by overexpression of gamma-8 (red). Wild-type CA1 pyramidal cells with a strong pedestal response (orange) tended to have larger miniature currents.

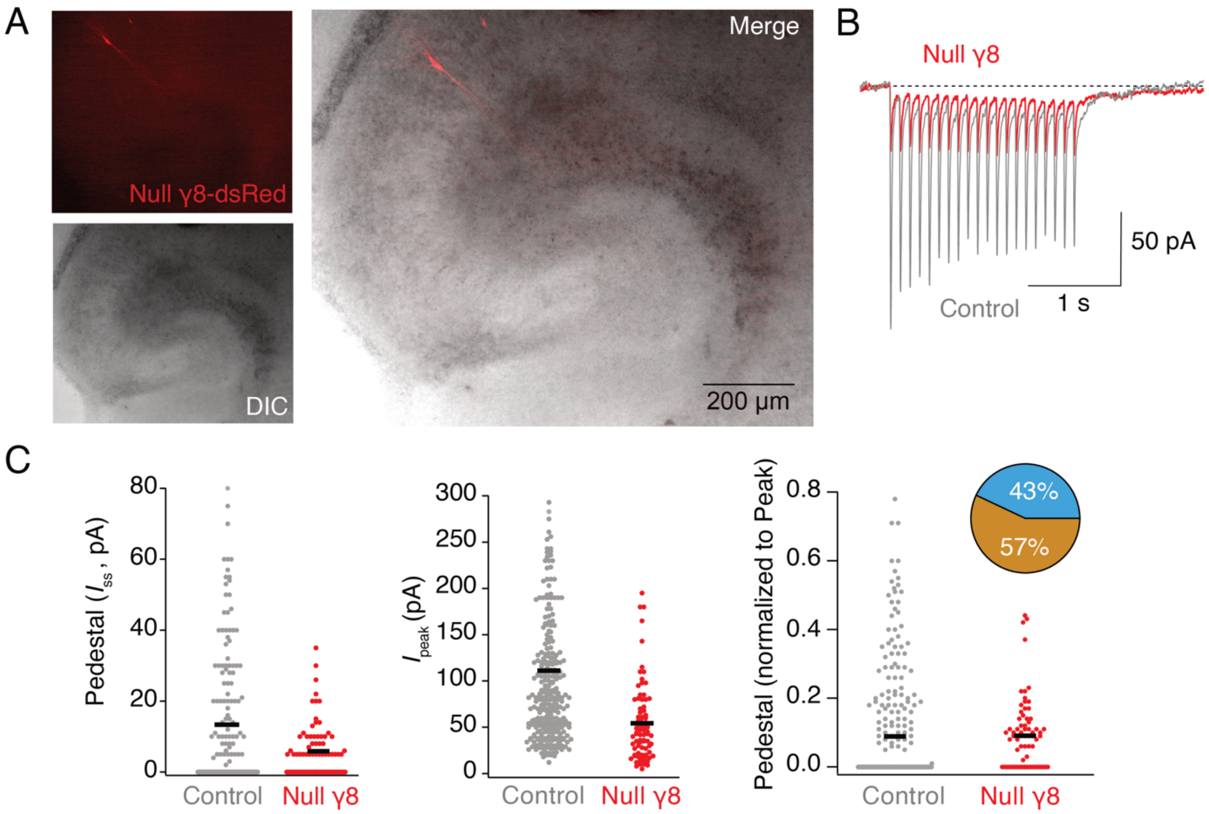


**Supplementary Figure 9. Null Gamma-8 electroporation and uncaging**

(A) Micrograph shows electroporation and expression of the null gamma-8 subunit from a vector also encoding dsRed in organotypic hippocampal slice. (B) Example response to 10 Hz uncaging from the neuron shown in panel (A). (C) On average, null gamma-8 reduced the pedestal amplitude but also the peak current amplitude. The mean amplitude of the normalized pedestal amplitude was approximately the same, but the incidence of the pedestal was increased (to 57%), and no large pedestal currents were observed. The average amplitude of the normalized pedestal (where observed, excluding the non-pedestal responses) was reduced from 26% in control to 14% in the presence of null gamma-8.
